# Optimized functional annotation of ChIP-seq data

**DOI:** 10.1101/082347

**Authors:** Bohdan B. Khomtchouk, William C. Koehler, Derek J. Van Booven, Claes Wahlestedt

## Abstract

**Motivation:** Different ChIP-seq peak callers often produce different output results from the same input. Since different peak callers are known to produce differentially enriched peaks with a large variance in peak length distribution and total peak count, accurately annotating peak lists with their nearest genes can be an arduous process. Functional genomic annotation of histone modification ChIP-seq data can be a particularly challenging task, as chromatin marks that have inherently broad peaks with a diffuse range of signal enrichment (e.g., H3K9me1, H3K27me3) differ significantly from narrow peaks that exhibit a compact and localized enrichment pattern (e.g., H3K4me3, H3K9ac). In addition, varying degrees of tissue-dependent broadness of an epigenetic mark can make it difficult to accurately and reliably link sequencing data to biological function. Thus, there exists an unmet need to develop a software program that can precisely tailor the computational analysis of a ChIP-seq dataset to the specific peak coordinates of the data and its surrounding genomic features.

**Results:** *geneXtendeR* optimizes the functional annotation of ChIP-seq peaks by exploring relative differences in annotating ChIP-seq peak sets to variable-length gene bodies. In contrast to prior techniques, *geneXtendeR* considers peak annotations beyond just the closest gene, allowing users to investigate peak summary statistics for the first-closest gene, second-closest gene, …, *n*^*th*^-closest gene whilst ranking the output according to biologically relevant events and iteratively comparing the fidelity of peak-to-gene overlap across a user-defined range of upstream and downstream extensions on the original boundaries of each gene’s coordinates. We tested *geneXtendeR* on 547 human transcription factor ChIP-seq ENCODE datasets and 198 human histone modification ChIP-seq ENCODE datasets, providing the analysis results as case studies.

**Availability:** The *geneXtendeR* R/Bioconductor package (including detailed introductory vignettes) is available under the GPL-3 Open Source license and is freely available to download from Bioconductor at: https://bioconductor.org/packages/geneXtendeR/

**Author summary:** *geneXtendeR* makes functional annotation of ChIP-seq data more robust and precise, regardless of peak variability attributable to parameter tuning or peak caller algorithmic differences. Since different ChIP-seq peak callers produce differentially enriched peaks with large variance in peak length distribution and total peak count, annotating peak lists with their nearest genes can often be a noisy process where an adjacent second or third-closest gene may constitute a more viable biological candidate, e.g., during cases of linked genes that are located close to each other. As such, the goal of *geneXtendeR* is to robustly link differentially enriched peaks with their respective genes, thereby aiding experimental follow-up and validation in designing primers for a set of prospective gene candidates during qPCR.

## Introduction

The field of epigenetic research studies the process by which heritable changes in gene expression occur without underlying alterations in the DNA sequence. Epigenetics plays a key role in human biology, and dysregulation in epigenetic processes is associated with the pathogenesis of cancer and many other diseases. Epigenetic mechanisms have been demonstrated to be necessary for biological programs important for a variety of health and disease outcomes. Understanding the impact of epigenetic architecture on the accessibility of gene promoters and its effect on gene expression patterns is therefore critical for linking chromatin biology to clinical indications. One way to measure such events involves investigating histone modifications, namely post-translational modifications to histones (referred to as chromatin marks) that regulate gene expression by organizing the genome into active regions of euchromatin, where DNA is accessible for transcription, or inactive heterochromatin regions, where DNA is more compact and less accessible for transcription [1].

Chromatin marks come in a variety of different shapes and sizes, ranging from the extremely broad to the extremely narrow [2–6]. This spectrum depends on a number of biological factors ranging from qualitative characteristics such as tissue-type [7] to temporal aspects such as developmental stage [8]. Depending on the peak caller used, computational factors such as the variance observed in peak coordinate positions (peak start, peak end) – both in terms of length distribution of peaks as well as the total number of peaks called – is an issue that persists even when samples are run at identical default parameter values [9, 10]. This variance becomes a factor when annotating peak lists genome-wide with their nearest genes as peaks can be shifted in genomic position (towards 5’ or 3’ end) or be of different lengths, depending on the peak caller employed. In total, the combined effect of these factors exerts a unique influence over the functional annotation and understanding of genomic variability, which ultimately complicates the study of epigenetic regulation of biological function.

Prior software in the ChIP-seq functional annotation arena (e.g., ANNOVAR [11], ChIPpeakAnno [12], ChIPseeker [13], HOMER [14], and BEDTools [15]) has focused exclusively on distance-minimizing algorithms between peaks and the transcriptional start site (TSS) regions of their nearest genes. In contrast, *geneXtendeR* significantly expands this definition to include n-dimensional annotation, whereby a user can investigate second-closest, third-closest, …, *n*^*th*^-closest genes to any given peak (or set of peaks), thereby focusing on and prioritizing the biology over simply the raw numbers (in base pairs). Detailed expositions of these new methods and their implications on the interpretation of results from data analyses are presented as case studies in the *geneXtendeR* package vignette.

## Materials and methods

### Algorithms and implementation

The key algorithm in the *geneXtendeR* R/Bioconductor package is the extension algorithm, implemented in the C programming language for performance and efficiency. The process of “extending” refers to performing sequential iterative gene-feature overlaps after adding to the gene-span a user-specified region upstream of the start of the gene model and a fixed (500 bp) region downstream of the gene, resulting in assigning to a gene the features that do not physically overlap with it but are sufficiently close. This process is repeated multiple times across a range of extension parameters set by the user and a series of visualizations are returned as output to help users hone in on the optimal functional annotation. This is in contrast to most past and present epigenetic analyses, in both ChIP-seq [16] and ATAC-seq [17] studies, that ad-hoc assign gene body definitions (e.g., assigning a default 2 kbp as the cutoff for gene-proximal peaks) before mapping the peaks to genomic features. Fig. 2 shows why such a practice may be limiting.

**Fig 1.**
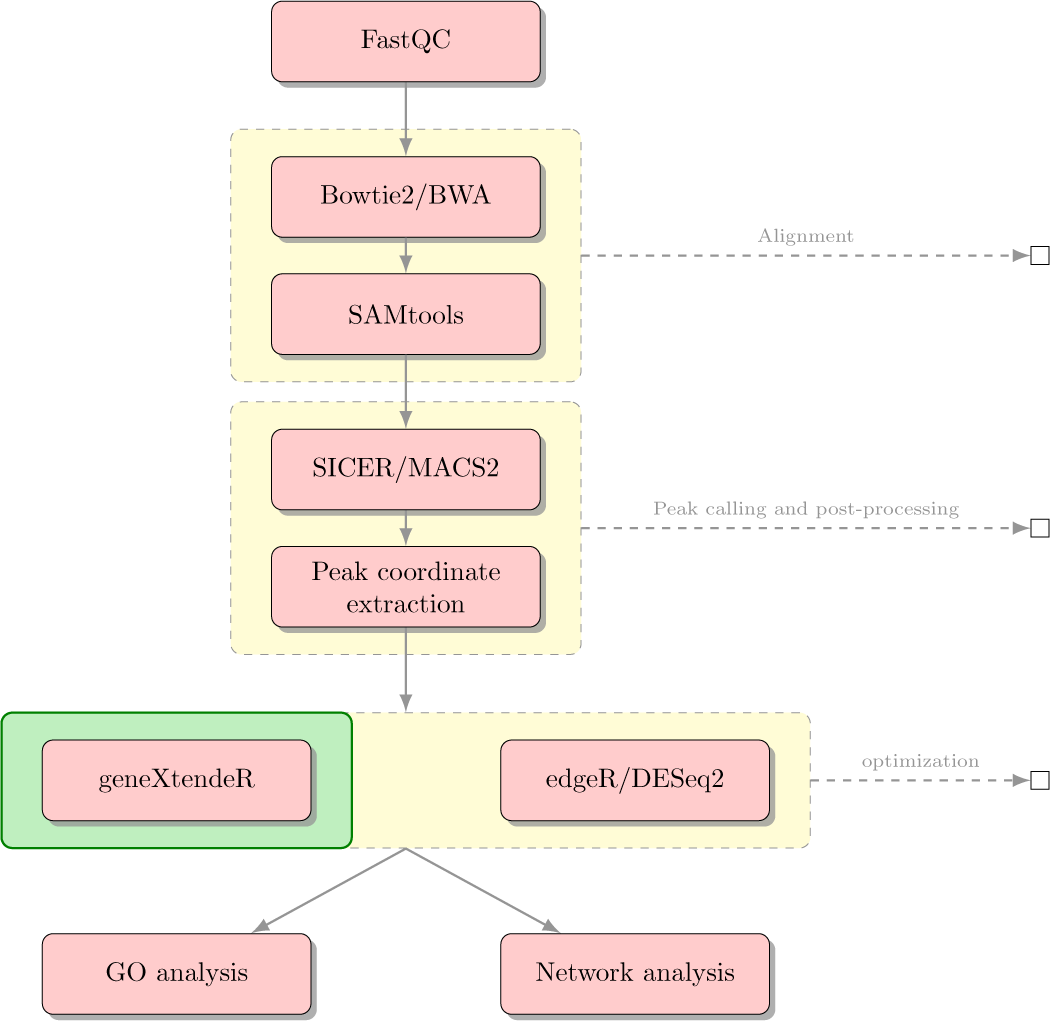
Sample biological workflow. Sample biological workflow using *geneXtendeR* in combination with existing statistical software to evaluate the role of ChIP-seq peak significance during functional annotation tasks (see description of hotspotPlot() function in package vignette). It is not uncommon for significant peaks to be located thousands of base pairs away from their nearest genes, suggesting that sequences under these respective peaks may further be extracted and analyzed for the presence of known regulatory elements or repeats (e.g., using software programs like TRANSFAC, MEME/JASPAR, or RepeatMasker) or for investigating potential enhancer effects.

**Fig 2.**
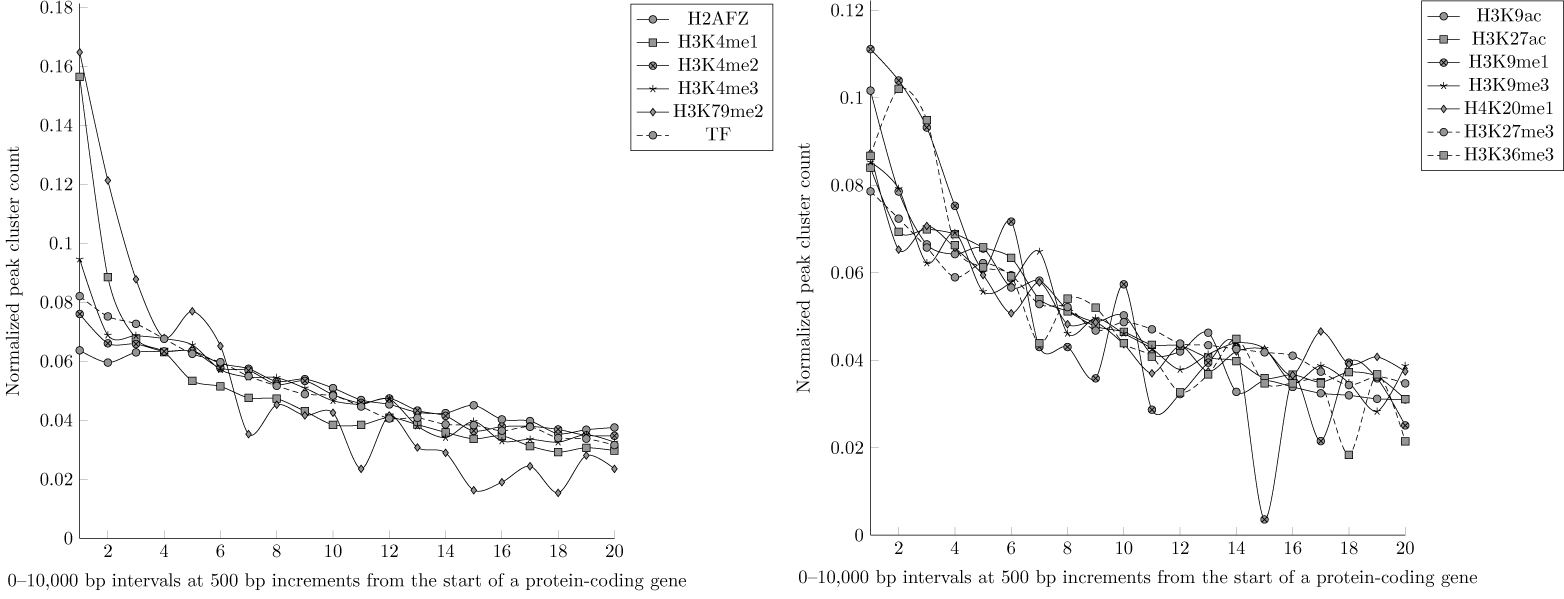
ENCODE ChIP-seq datasets. Large-scale computational *geneXtendeR* analysis using hg19 reference genome of 198 histone modification and 547 transcription factor ChIP-seq datasets from ENCODE. To make data directly comparable to each other, the y-axis represents a normalized count of peak clusters (number of peak clusters in a specific interval divided by the total number of peak clusters across all 0-10 kbp intervals for a given chromatin mark or TF), where a peak cluster is defined as a genomic locus harboring at least 5 overlapping peaks. The x-axis, which is segmented into 20 discrete regions (“1” = 0-500 bp interval, “2” = 500-1000 bp interval, …, “20” = 9500-10000 bp interval), represents a genomic distance (in bp) of the closest protein-coding gene to each respective peak cluster. A steady decline in peak cluster count at further upstream intervals is detected for all (broad and narrow) chromatin marks as well as transcription factors, i.e., peak clusters do not congregate proximally within any specific region of intervals (e.g., 0-2000 bp) of their respective protein-coding genes, as there is a large number of peak clusters that reside further upstream of their nearest gene. For instance, in the 9500-10000 bp interval alone, there are 1043 peak clusters for the H2AFZ chromatin mark, 569 peak clusters for the H3K4me1 chromatin mark, and 716 peak clusters across all transcription factor ChIP-seq datasets. However, there are certainly exceptions like the H3K9me1 chromatin mark, which has only 1 peak cluster in the 7000-7500 bp interval (see the big dip at x-axis=15 in the right-hand panel) and only 7 peak clusters in the 9500-10000 bp interval (see S1 Appendix and S2 Appendix for reproducible code and data).

From a performance standpoint, the extension algorithm is optimized to handle the computational complexity inherent to performing compute-intensive n-dimensional annotation. This ultimately aids in efficiently capturing cis-regulatory and proximal-promoter element relationships between ChIP-seq peaks and the genes they are (dys-)regulating, as described in further detail in the vignette. All of *geneXtendeR*’s source code is implemented in the C and R programming languages and shipped within a standalone R/Bioconductor package release that is publicly available for download from either Bioconductor or Github. Within its codebase, *geneXtendeR* leverages the AnnotationDbi [18], BiocStyle [19], data.table [20], dplyr [21], GO.db [22], networkD3 [23], RColorBrewer [24], rtracklayer [25], SnowballC [26], testthat [27], tm [28], and wordcloud [29] libraries.

### Biological Workflow

Fig 1 summarizes the key steps of a sample workflow. For an end-to-end example of a comprehensive biological workflow and case-study, please refer to the vignette.

## Results

First, we tested *geneXtendeR* on all publicly available transcription factor and histone modification ChIP-seq datasets in ENCODE. After downloading and analyzing data from the ENCODE ChIP-seq Experiment Matrix (hg19) [30], our large-scale analysis (Fig. 2) indicated that ChIP-seq peaks do not concentrate within any specific upstream extension (e.g., 2000 bp) of their nearest protein-coding genes. This observation that ChIP-seq peaks drop off gradually with genomic distance from the start of a gene (first exon) suggests that there is no good general guideline cutoff for capturing proximal histone modifications (e.g., prior studies [16, 17] have used 2000 bp) or transcription factor binding peaks. There are still hundreds of peak clusters that reside in proximal-promoter regions that are 2000-3000 bp away from their nearest protein-coding genes and in distal regions beyond 3 kbp, making ad-hoc decisions like 2 kbp cutoffs too general to be of broad utility across specific use cases. When applying *geneXtendeR* to both proximal and distal transcription factor (TF) binding peaks for all cell types, we observed some cell type-dependent and TF-dependent peak aggregation dynamics in intervals ranging from 0 to 10 kbp (Fig 3). Similarly, examining distal peaks in representative plots of different chromatin marks in different cell types indicated that peaks indeed aggregate in a cell type and chromatin mark-dependent manner (Fig 4). S1 Appendix and S2 Appendix provide downloads to the complete compendium of all proximal/distal datasets analyzed from ENCODE.

**Fig 3.**
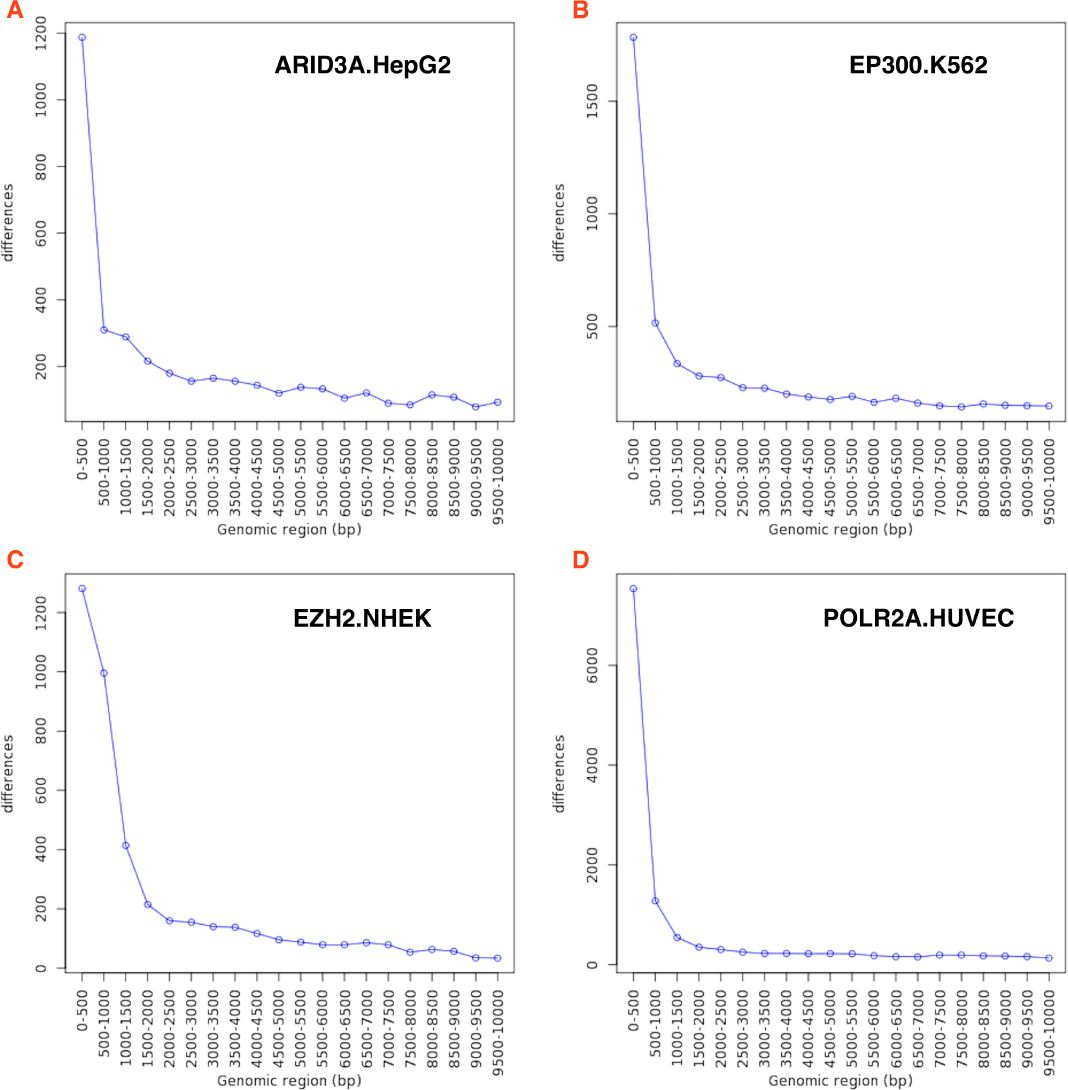
ENCODE TF analysis. Running *geneXtendeR* on 547 human transcription factor (TF) ChIP-seq datasets obtained from ENCODE shows that many peaks tend to reside within 500 bp upstream of their respective protein-coding genes yet, depending on the identity of the transcription factor (e.g., EP300) and the specific cell type (e.g., K562), there may be more or less peaks located further upstream and, therefore, a generalized upstream cutoff is not applicable.

**Fig 4.**
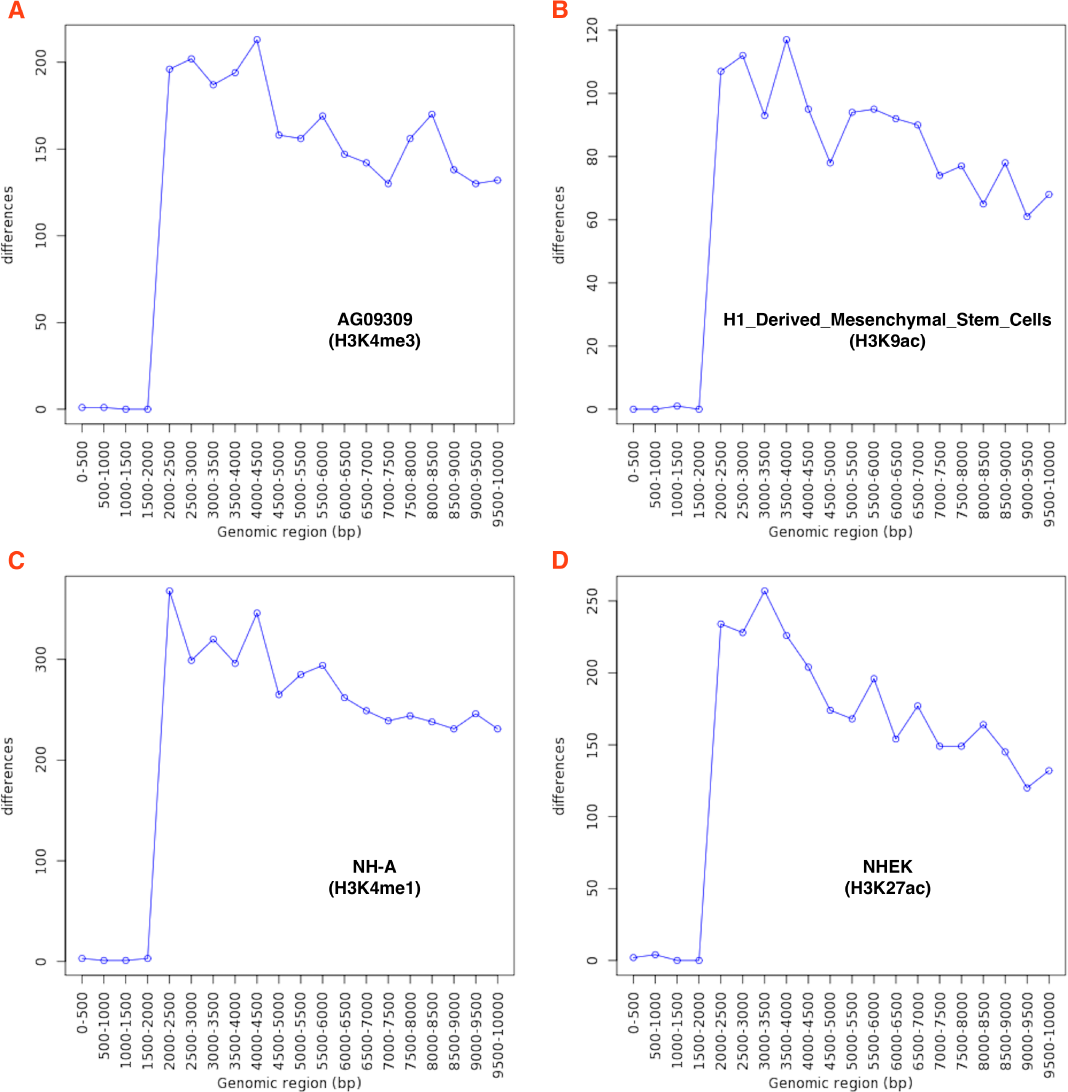
ENCODE histone modification analysis. Running *geneXtendeR* on 198 human histone modification ChIP-seq distal peak datasets obtained from ENCODE reveals that most distal peaks are not congregating within any specific upstream region of their respective protein-coding genes (here we define “distal” as only those peaks that are more than 2000 bp away from their nearest gene). Additional comprehensive analyses (see S1 Appendix and S2 Appendix) were run for proximal peaks (≤ 2000 bp) as well as the complete set of peaks (proximal + distal) from all 198 histone modification ChIP-seq datasets, and similar patterns were observed.

We then focused our attention on using *geneXtendeR* to perform an end-to-end analysis of a published histone modification ChIP-seq dataset [31] deposited in the Gene Expression Omnibus under accession number GSE83979. At the peak-calling stage (Fig 1) we ran two different peak callers (SICER [32] and CisGenome [33]) producing two highly variable peak length profiles even at default run parameters S1 Fig. Despite the stark difference in peak profiles, *geneXtendeR* consistently identified the same top two gene candidates, highlighting its utility for robust functional annotation even in the face of extreme peak variability. Details are discussed in the package vignette.

We followed up this computational analysis by performing *n*-dimensional annotation of the GSE83979 dataset to provide an expanded view of the gene neighborhood around each individual peak – effectively annotating every peak *n* times (once for the closest gene, once for the second-closest gene, etc.) and grouping the results into a tabular summary format. We show in the vignette how the second-closest gene may be a preferable candidate for experimental follow-up/validation, especially if the first-closest gene is putative/predicted, while the second-closest gene is known to play a role in a similar biological process based on previously published literature.

## Discussion

The cell-type and TF/chromatin mark-specific complexity apparent in Fig. 3 and Fig. 4 motivated the design and implementation of user-friendly functions that can calculate ratios of statistically significant peaks to total peaks in various genomic intervals (see hotspotPlot() documentation in *geneXtendeR* vignette). Similarly, users can transform peaks into merged peaks (see peaksMerge()). *geneXtendeR* also allows users to explore gene ontology differences at various extensions (see diffGO()) as interactive network graphics (see makeNetwork()) or word clouds (see makeWordCloud()). Furthermore, users can investigate mean (average) peak lengths within any genomic interval (see meanPeakLengthPlot()), showing how average peak broadness can change at different upstream extensions, or examine the variance of peak lengths within a specific genomic interval (see peakLengthBoxplot()). It is also possible to examine unique genes and their associated ChIP-seq peaks between any two upstream extension levels (see distinct()). For example, Fig. 5 displays all unique genes (and their respective gene ontologies) that are associated with peaks located between 2-3 kbp across the genome. *geneXtendeR* also allows users to examine the distribution of peak lengths across the entire peak set (see allPeakLengths()), a function that is useful for visualizing the length distribution of all peaks from a peak caller. These functions (and more) are all explored in detail within the package vignette. After a user has explored the peak coordinates data using these functions to determine the optimal alignment of peaks to a GTF file, the peaks file can be functionally annotated with the annotate() function or one of its counterparts (gene annotate() or annotate n()) for n-dimensional annotation.

**Fig 5.**
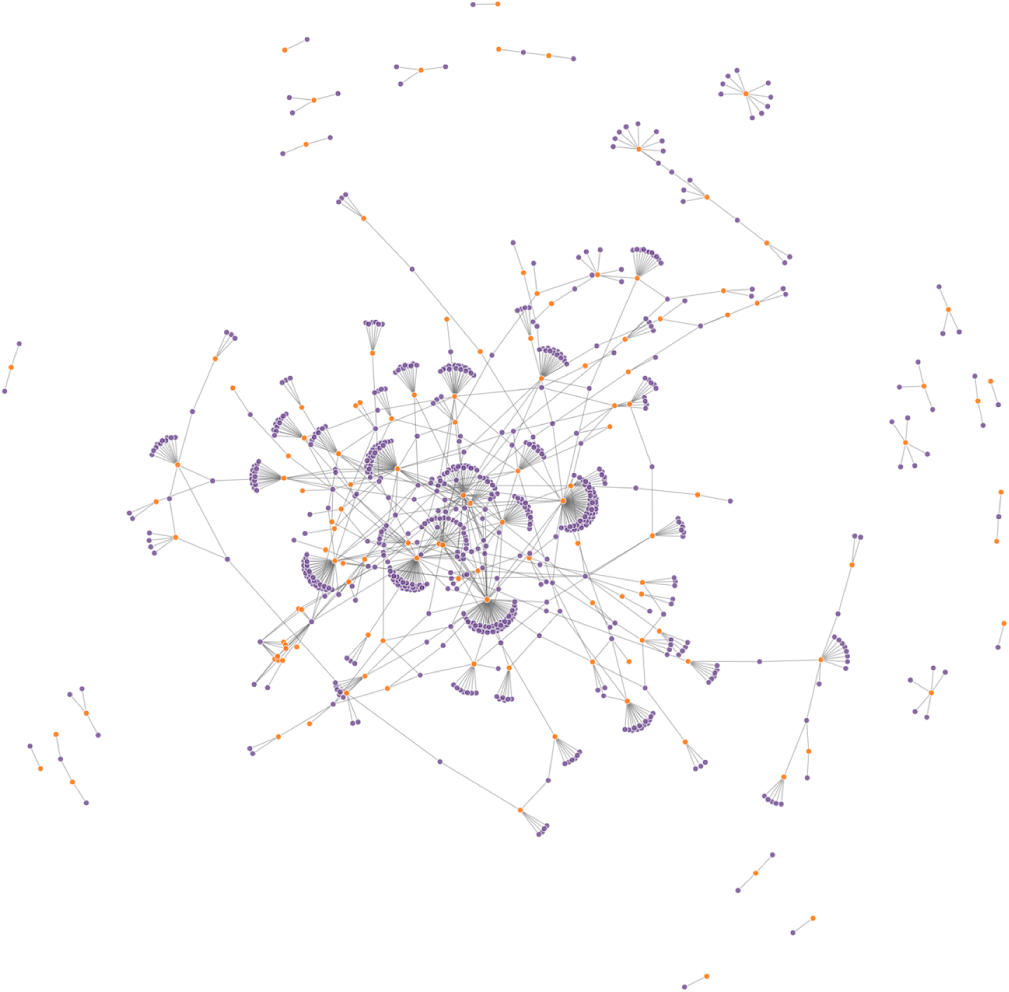
Genome-wide network analysis of peak subsets in promoter regions. All unique genes (and their respective gene ontologies (GO)) that are associated with peaks located in promoter-proximal regions between 2-3 kbp genome-wide. Put another way, these are all gene-GO pairs associated with peaks that are distinct between 2000 and 3000 bp upstream extensions across the genome. Orange color denotes gene names, purple color denotes GO terms. A user can hover the mouse cursor over any given node to display its respective label directly within RStudio. Likewise, users can dynamically drag and re-organize the spatial orientation of nodes, as well as zoom-in and out of them for visual clarity.

We have successfully applied *geneXtendeR* during the analysis of a histone modification ChIP-seq study investigating the neuroepigenetics of alcohol addiction [34], where *geneXtendeR* was used to determine an optimal upstream extension cutoff for H3K9me1 enrichment (a commonly studied broad peak) in rat brain tissue based on line plots of both significant peaks and total peaks. This analysis helped us to identify, functionally annotate, and experimentally validate synaptotagmin 1 (Syt1) as a key mediator in alcohol addiction and dependence [34]. This analysis is explored in detail in the package vignette. Taken together, *geneXtendeR*’s functions are designed to be used as an integral part of a broader biological workflow (Fig. 1).

## Conclusion

We present an R/Bioconductor package, *geneXtendeR*, that goes beyond the typical nearest-to-gene analyses commonplace to most standard computational ChIP-seq workflows. *geneXtendeR* offers n-dimensional functional annotation and the ability to investigate the effect of variable-length gene bodies when mapping peaks to genomic features, thereby serving as a next-generation model of peak annotation to nearby features in modern bioinformatics workflows. *geneXtendeR* therefore represents a critical first step towards tailoring the functional annotation of a ChIP-seq peak dataset according to the details of the peak coordinates (chromosome number, peak start position, peak end position) and their surrounding genomic features.

## Supporting information

**S1 Fig.**
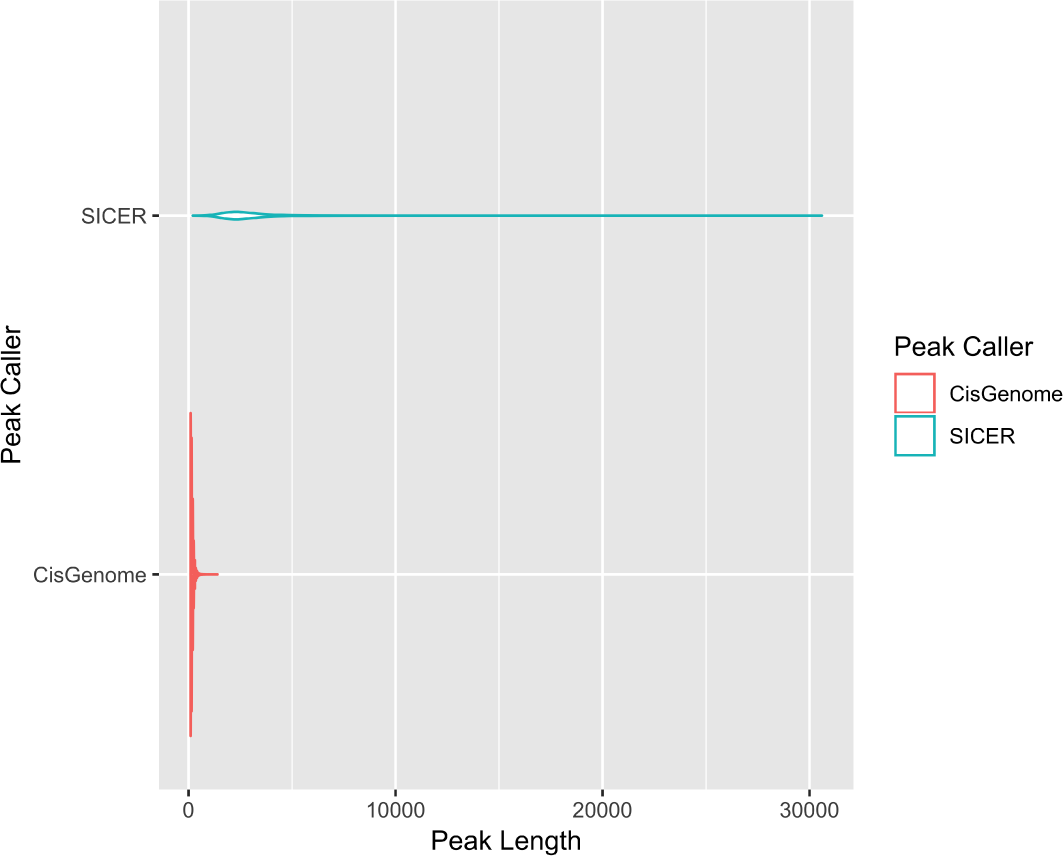
SICER vs. CisGenome peak length distribution differences for GSE83979. Violin plot showing the differences in peak length distributions of the same ChIP-seq data (available through the Gene Expression Omnibus database, accession identifier GSE83979) analyzed with two separate peak callers (SICER and CisGenome) – despite significant differences in peak lengths generated by the two callers (i.e., peak variability), *geneXtendeR*’s gene annotate() function can still robustly call top gene candidates consistently, as explained in the *geneXtendeR* package vignette.

**S1 Appendix. *geneXtendeR* analysis on 547 human TF ChIP-seq ENCODE datasets.** Files available here: https://github.com/Bohdan-Khomtchouk/ENCODE_TF_geneXtendeR_analysis

**S2 Appendix. *geneXtendeR* analysis on 198 human histone modification ChIP-seq ENCODE datasets.** Files available here: https://github.com/Bohdan-Khomtchouk/ENCODE_histone_geneXtendeR_analysis

### Acknowledgments

This work was supported by the American Heart Association (AHA) Postdoctoral Fellowship grant #18POST34030375 (Khomtchouk). This work was also partially supported by the Stanford Training Program in Aging Research grant (NIH/NIA T32-AG0047126) and the Army Research Office (ARO), National Defense Science and Engineering Graduate (NDSEG) Fellowship, 32 CFR 168a – both awarded to BBK from 2014-2018. The content is solely the responsibility of the authors and does not necessarily represent the official views of the American Heart Association, National Institutes of Health, or Department of Defense.

### Disclosures

BBK is a co-founder of Quiltomics.

## References

1. Abcam. Histone modifications: a guide. https://www.abcam.com/epigenetics/histone-modifications-a-guide

2. Squazzo SL, O’Geen H, Komashko VM, Krig SR, et al. Suz12 binds to silenced regions of the genome in a cell-type-specific manner. Genome Research. 2006, 16: 890–900.

3. Pepke S, Wold B, Mortazavi A, et al. Computation for ChIP-seq and RNA-seq studies. Nature Methods. 2009, 6 (11 Suppl): S22–S32.

4. Landt SG, Marinov GK, Kundaje A, Kheradpour P, et al. ChIP-seq guidelines and practices of the ENCODE and modENCODE consortia. Genome Research. 2012, 22(9): 1813–1831.

5. Kellis M, Wold B, Snyder MP, Bernstein BE, et al. Defining functional DNA elements in the human genome. Proceedings of the National Academy of Sciences. 2014, 111(17): 6131–6138.

6. Heinig M, Colomé-Tatché M, Taudt A, Rintisch C, et al. histoneHMM: Differential analysis of histone modifications with broad genomic footprints. BMC Bioinformatics. 2015, 16:60.

7. Rintisch C, Heinig M, Bauerfeind A, Schafer S, et al. Natural variation of histone modification and its impact on gene expression in the rat genome. Genome Research. 2014, 24(6): 942–953.

8. Ha M, Ng DW, Li WH, Chen ZJ. Coordinated histone modifications are associated with gene expression variation within and between species. Genome Research. 2011, 21(4): 590–598.

9. Koohy H, Down TA, Spivakov M, Hubbard T. A Comparison of Peak Callers Used for DNase-Seq Data. PLoS One. 2014, 9(8): e105136.

10. Thomas R, Thomas S, Holloway AK, Pollard KS. Features that define the best ChIP-seq peak calling algorithms. Briefings in Bioinformatics. 2017, 18(3): 441–450.

11. Wang K, Li M, Hakonarson H. ANNOVAR: functional annotation of genetic variants from high-throughput sequencing data. Nucleic Acids Research. 2010, 38(16): e164.

12. Zhu L, Gazin C, Lawson N, Pagès H, et al. ChIPpeakAnno: a Bioconductor package to annotate ChIP-seq and ChIP-chip data. BMC Bioinformatics. 2010, 11(1), pp. 237.

13. Yu G, Wang LG, He QY. ChIPseeker: an R/Bioconductor package for ChIP peak annotation, comparison and visualization. Bioinformatics. 2015, 31(14): 2382–2383.

14. Heinz S, Benner C, Spann N, Bertolino E, et al. Simple Combinations of Lineage-Determining Transcription Factors Prime cis-Regulatory Elements Required for Macrophage and B Cell Identities. Mol Cell 2010, 38(4): 576–589.

15. Quinlan AR, Hall IM. BEDTools: a flexible suite of utilities for comparing genomic features. Bioinformatics. 2010, 26(6): 841–842.

16. Maze I, Feng J, Wilkinson MB, Sun H, et al. Cocaine dynamically regulates heterochromatin and repetitive element unsilencing in nucleus accumbens. Proceedings of the National Academy of Sciences. 2011, 108(7): 3035–3040.

17. Wang J, Zibetti C, Shang P, Sripathi SR, et al. ATAC-Seq analysis reveals a widespread decrease of chromatin accessibility in age-related macular degeneration. Nature Communications. 2018, 9:1364.

18. Pagès H, Carlson M, Falcon S, Li N. AnnotationDbi: Annotation Database Interface. R package version 1.42.1 (2018).

19. Oleś A, Morgan M, Huber W. BiocStyle: Standard styles for vignettes and other Bioconductor documents. R package version 2.8.2 (2018).

20. Dowle M, Srinivasan A. data.table: Extension of ‘data.frame‘. R package version 1.11.4 (2018).

21. Wickham H, François R, Henry L, Müller K. dplyr: A Grammar of Data Manipulation. R package version 0.7.6 (2018).

22. Carlson M. GO.db: A set of annotation maps describing the entire Gene Ontology. R package version 3.6.0 (2018).

23. Allaire JJ, Gandrud C, Russell K, Yetman CJ. networkD3: D3 JavaScript Network Graphs from R. R package version 0.4 (2017).

24. Neuwirth E. RColorBrewer: ColorBrewer Palettes. R package version 1.1.2 (2014).

25. Lawrence M, Gentleman R, Carey V. rtracklayer: an R package for interfacing with genome browsers. Bioinformatics. 2009, 25: 1841–1842.

26. Bouchet-Valat M. SnowballC: Snowball stemmers based on the C libstemmer UTF-8 library. R package version 0.5.1 (2014).

27. Wickham H. testthat: Get Started with Testing. The R Journal. 2011, 3(1): 5–10.

28. Feinerer I, Hornik K, Meyer D. Text Mining Infrastructure in R. Journal of Statistical Software. 2008, 25(5): 1–54.

29. Fellows I. wordcloud: Word Clouds. R package version 2.5 (2014).

30. https://genome.ucsc.edu/encode/dataMatrix/encodeChipMatrixHuman.html

31. Gidlöf O, Johnstone AL, Bader K, Khomtchouk BB, et al. Ischemic Preconditioning Confers Epigenetic Repression of Mtor and Induction of Autophagy Through G9a-Dependent H3K9 Dimethylation. Journal of the American Heart Association: Cardiovascular and Cere-brovascular Disease. 2016, 5(12): e004076.

32. Zang C, Schones DE, Zeng C, Cui K, et al. A clustering approach for identification of enriched domains from histone modification ChIP-Seq data. Bioinformatics. 2009, 25(15): 1952–1958.

33. Ji H, Jiang H, Ma W, Johnson DS, et al. An integrated software system for analyzing ChIP-chip and ChIP-seq data. Nature Biotechnology. 2008, 26: 1293–1300.

34. Barbier E, Johnstone AL, Khomtchouk BB, Tapocik JD, et al. Dependence-induced increase of alcohol self-administration and compulsive drinking mediated by the histone methyltransferase PRDM2. Molecular Psychiatry. 2017, 22: 1746–1758.

